# Biochemical evidence for the diversity of LHCI proteins in PSI-LHCI from the red alga *Galdieria sulphuraria* NIES-3638

**DOI:** 10.1101/2024.11.04.622003

**Authors:** Ryo Nagao, Haruya Ogawa, Takehiro Suzuki, Naoshi Dohmae, Koji Kato, Yoshiki Nakajima, Jian-Ren Shen

**Affiliations:** Faculty of Agriculture, Shizuoka University, Shizuoka 422-8529, Japan; Research Institute for Interdisciplinary Science and Graduate School of Natural Science and Technology, Okayama University, Okayama 700-8530, Japan; Biomolecular Characterization Unit, RIKEN Center for Sustainable Resource Science, Saitama 351-0198, Japan

**Author notes:** Corresponding Author: Ryo Nagao. Abbreviations: *β*-DDM, *n*-dodecyl-*β*-D-maltoside; Car, carotenoid; Chl, chlorophyll; LHC, light-harvesting complex; LHCI, PSI-specific LHCs; PSI, photosystem I.

**Keywords:** *Galdieria sulphuraria* NIES-3638, PSI-LHCI, Red alga, RedCAP

## Abstract

Red algae are photosynthetic eukaryotes whose light-harvesting complexes (LHCs) associate with photosystem I (PSI). In this study, we examined characteristics of PSI-LHCI, PSI, and LHCI isolated from the red alga *Galdieria sulphuraria* NIES-3638. The PSI-LHCI supercomplexes were purified using anion-exchange chromatography followed by hydrophobic interaction chromatography, and finally by trehalose density gradient centrifugation. PSI and LHCI were similarly prepared following the dissociation of PSI-LHCI with Anzergent 3-16. Polypeptide analysis of PSI-LHCI revealed the presence of PSI and LHC proteins, along with a red-lineage chlorophyll *a*/*b*-binding-like protein (RedCAP), which is distinct from LHC proteins within the LHC protein superfamily. RedCAP, rather than LHC proteins, exhibited tight binding to PSI. Carotenoid analysis of LHCI identified zeaxanthin, *β*-cryptoxanthin, and *β*-carotene, with zeaxanthin particularly enriched, which is consistent with other red algal LHCIs. A Qy peak of chlorophyll *a* in the LHCI absorption spectrum was blue-shifted compared with those of PSI-LHCI and PSI, and a fluorescence emission peak was similarly shifted to shorter wavelengths. Based on these results, we discuss the diversity of LHC proteins, including RedCAP, in red algal PSI-LHCI supercomplexes.

## Introduction

In cyanobacteria, algae, and land plants, oxygenic photosynthesis converts solar energy into chemically usable energy (Blankenship 2021). This process is driven by multi-subunit protein complexes, known as photosystem I (PSI) and photosystem II (PSII) (Blankenship 2021). PSI facilitates light-induced electron transfer from plastocyanin or cytochrome c6 on the lumenal side of the thylakoid membrane to ferredoxin on the stromal side (or the cytosolic side in cyanobacteria) (Brettel and Leibl 2001; Fromme et al. 2001; Golbeck 1992; Nelson and Junge 2015). To support this reaction, light-harvesting antenna complexes are associated with PSI, transferring the absorbed energy to the photosystem (Blankenship 2021).

Photosynthetic organisms have evolved a diverse array of light-harvesting antennae over the course of evolution (Blankenship 2021). These antennae are classified into two main groups: membrane proteins and water-soluble proteins. Distinct antenna types contribute to the color diversity observed among photosynthetic organisms (Falkowski et al. 2004). A key group within the membrane protein category is the light-harvesting complex (LHC) protein superfamily (Engelken et al. 2010; Sturm et al. 2013). In photosynthetic eukaryotes, PSI-specific LHCs (LHCIs) bind to a PSI monomer, forming a PSI-LHCI supercomplex. Notably, the binding properties of LHCIs within these supercomplexes exhibit substantial variability across different photosynthetic species (Hippler and Nelson 2021; Shen 2022).

Red algae represent a distinct photosynthetic lineage that encompasses both unicellular and large multicellular forms (Yoon et al. 2010). Previous studies have successfully isolated and characterized PSI-LHCI supercomplexes from various red algal species, including *Porphyridium cruentum*, *Galdieria sulphuraria*, *Cyanidium caldarium*, and *Cyanidioschyzon merolae* (Busch et al. 2010; Gardian et al. 2007; Haniewicz et al. 2018; Marquardt and Rhiel 1997; Nagao et al. 2023; Thangaraj et al. 2011; Tian et al. 2017; Wolfe et al. 1994). Among these, *G. sulphuraria*, a member of Cyanidiophyceae (Liu et al. 2020; Ott 2009; Ott and Seckbach 1994; Park et al. 2023), inhabits thermo-acidic environments (Gross and Schnarrenberger 1995). The structural and light-harvesting properties of PSI-LHCI supercomplexes from *G. sulphuraria* have been explored through negative-stain electron microscopy and time-resolved fluorescence spectroscopy (Thangaraj et al. 2011). However, the absorption and fluorescence characteristics, as well as the pigment composition of *G. sulphuraria* LHCIs, remain unexamined. This gap in knowledge is essential for understanding the light-harvesting and excitation-energy transfer mechanisms in the PSI-LHCI, PSI, and LHCI complexes of *G. sulphuraria*, as studied previously in other organisms (Giera et al. 2018; Nagao et al. 2019a; Nagao et al. 2023; Nagao et al. 2019c, d, 2020; Wientjes et al. 2011).

In this study, we purified three types of protein complexes, namely PSI-LHCI, PSI, and LHCI, from the red alga *G. sulphuraria* NIES-3638, and analyzed their protein and pigment compositions, as well as spectral characteristics. Biochemical and spectroscopic analyses unveiled distinct features of the *G. sulphuraria* PSI-LHCI supercomplex.

## Materials and methods

### Cell culture and thylakoid preparations

The red alga, *G. sulphuraria* NIES-3638, was grown in an inorganic medium (Allen 1959) at a photosynthetic photon flux density of 30 µmol photons m^−2^ s^−1^ at 30 °C with bubbling of air containing 3% (v/v) CO_2_. Following centrifugation, the cells were suspended with a 20 mM MES-NaOH (pH 6.5) buffer containing 0.2 M trehalose, 5 mM CaCl_2_, and 10 mM MgCl_2_ (buffer A). The harvested cells were disrupted by glass beads (Nagao et al. 2017). After removal of unbroken cells by centrifugation at 2,000 × g for 5 min at 4 °C, the resultant supernatant was centrifuged at 159,000 × g for 30 min, providing thylakoid membranes as resultant pellets. Thylakoid membranes were suspended in buffer A and stored at −80 °C.

### Preparation of PSI-LHCI supercomplexes

Isolation of PSI-LHCI was carried out at 4 °C unless otherwise indicated. Thylakoids were solubilized with 1% (w/v) *n*-dodecyl-*β*-D-maltoside (*β*-DDM) at a Chl concentration of 0.5 mg mL^−1^ by gentle agitation for 20 min on ice in the dark. Following removal of unsolubilized thylakoids by centrifugation at 162,000 × g for 20 min, the resultant supernatant was applied to a Q-Sepharose High Performance anion-exchange column (an inner diameter of 1.6 cm and a length of 25 cm) equilibrated with buffer A containing 0.03% *β*-DDM (buffer B). The column was washed with buffer B until the eluate became colorless. Elution was performed at a flow rate of 1.0 mL min^−1^ using linear gradients of buffer B containing 500 mM NaCl (buffer C): 0–600 min, 0–50% buffer C; 600–700 min, 100% buffer C. The fraction enriched in PSI-LHCI was eluted at 80–125 mM NaCl (labeled as 1 in Figure 1A).

**Figure 1.**
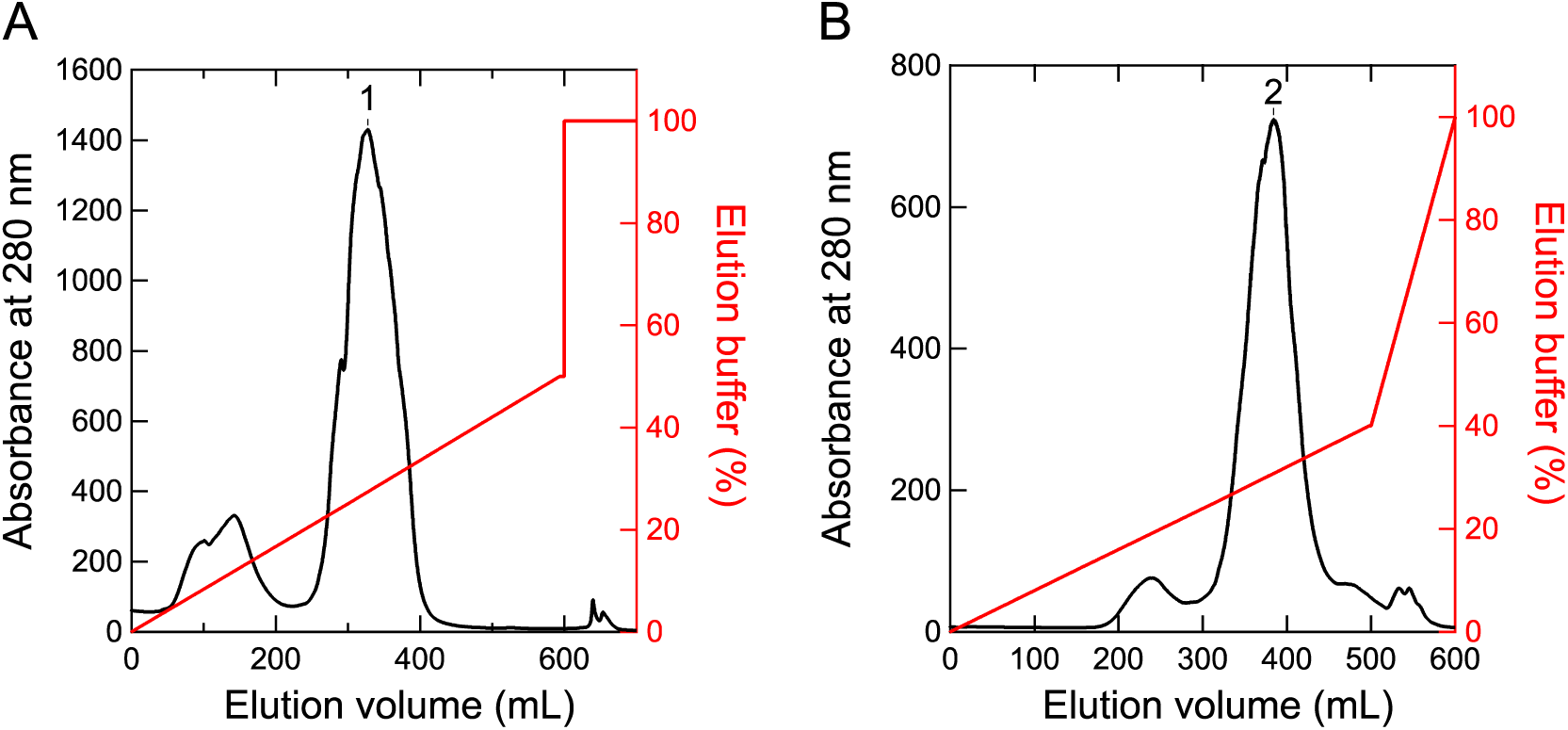
Fractionations of pigment-protein complexes. Elution profiles by anion-exchange chromatography (**A**) and hydrophobic-interaction chromatography (**B**). Fractions labeled as 1 and 2 were collected.

The PSI-LHCI enriched fraction was mixed with two volumes of a 20 mM MES-NaOH (pH 6.5) buffer containing 3 M ammonium sulfate and 0.03% *β*-DDM at a final concentration of 2 M ammonium sulfate. The mixed sample was applied to a Toyopearl Ether 650-M column (an inner diameter of 1.6 cm and a length of 12.5 cm) equilibrated with a 20 mM MES-NaOH (pH 6.5) buffer containing 2 M ammonium sulfate and 0.03% *β*-DDM (buffer D). The column was washed with buffer D until the eluate became colorless. Elution was performed at a flow rate of 1.0 mL min^−1^ using linear gradients of a 20 mM MES-NaOH (pH 6.5) buffer containing 0.03% *β*-DDM (buffer E): 0–500 min, 0–40% buffer E; and 500–600 min, 40–100% buffer E. The PSI-LHCI fraction was eluted at 1.53–1.34 M ammonium sulfate (labeled as 2 in Figure 1B), and centrifuged at 48,000 × g for 10 min after the addition of 50% (w/v) polyethylene glycol 1500 at a final concentration of 15%. The resultant precipitation was suspended with buffer A containing 0.03% *β*-DDM, and subsequently loaded onto a linear gradient of 10–40% trehalose in a medium containing 20 mM MES-NaOH (pH 6.5), 100 mM NaCl, 5 mM CaCl_2_, and 10 mM MgCl_2_, and 0.03% *β*-DDM. After centrifugation at 154,000 × g for 18 h (P40ST rotor; Hitachi), the purified PSI-LHCI supercomplexes were found in ∼25% trehalose layer, and concentrated using a 150 kDa cut-off filter (Apollo; Orbital Biosciences). The PSI-LHCI supercomplexes were stored at −80 °C.

### Preparation of PSI cores and LHCIs

The purified PSI-LHCI supercomplexes were treated with 2% *β*-DDM and 2% (w/v) Anzergent 3-16 at a Chl concentration of 0.5 mg mL^−1^ for 30 min at 4 °C in the dark, as described previously (Haworth et al. 1983; Nagao et al. 2023). The treated samples were loaded onto a linear gradient of 10–30% trehalose in a medium containing 20 mM MES-NaOH (pH 6.5), 100 mM NaCl, 5 mM CaCl_2_, and 10 mM MgCl_2_, and 0.03% *β*-DDM. After centrifugation at 154,000 × g for 18 h (P40ST rotor), each green band was collected (Figure 2A) and stored at −80 °C.

**Figure 2.**
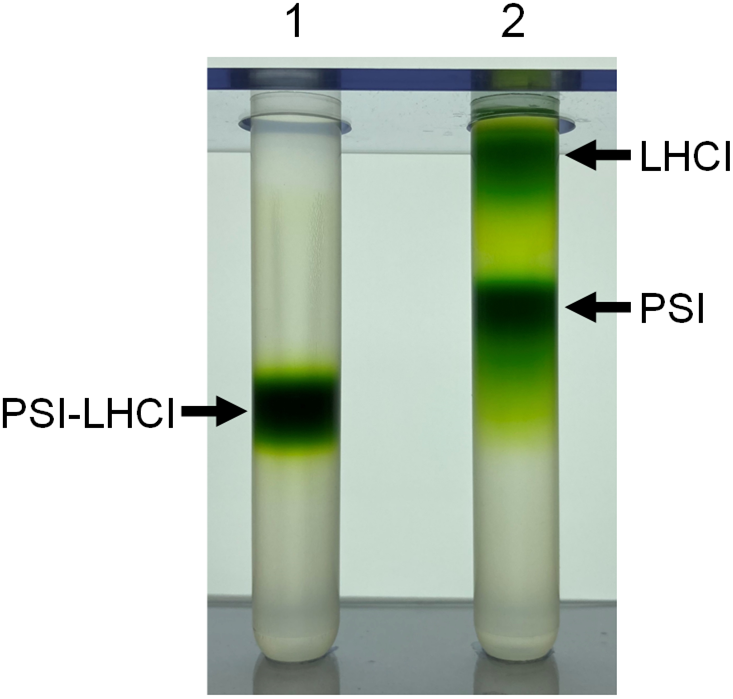
Profiles of trehalose density gradient centrifugation. Tube 1, untreated PSI-LHCI supercomplexes; tube 2, PSI-LHCI supercomplexes treated with Anzergent 3-16.

### SDS-PAGE and mass spectrometry

SDS-PAGE was performed according to the method of Ikeuchi and Inoue (1988). Three preparations of PSI-LHCI (5 µg of Chl), PSI (5 µg of Chl), and LHCI (2 µg of Chl), were incubated for 30 min after the addition of 3% lithium lauryl sulfate and 75 mM dithiothreitol to each sample and then subjected to SDS-PAGE using a 16% polyacrylamide gel containing 7.5 M urea. A standard molecular weight marker (SP-0110; APRO Science) was used. After electrophoresis, the SDS-PAGE gel was stained with Coomassie Brilliant Blue R-250. Mass spectrometry was performed according to our previous study (Nagao et al. 2019b).

### Pigment analysis

Pigment compositions were analyzed using a Shimadzu HPLC system equipped with a reversed-phase Inertsil C8 column (150 × 4.6 mm, 5 µm particle size; GL Sciences) (Nagao et al. 2019c). Pigments were extracted from each sample through the use of 100% methanol and eluted from the column employing solvent A (methanol:acetonitrile:0.25 M pyridine = 50:25:25 (v:v:v)) and solvent B (methanol:acetonitrile:acetone = 20:60:20 (v:v:v)) (Zapata et al. 2000). Elution was conducted in accordance with the method of Zapata et al. (2000). The flow rate was set to 0.9 mL min^−1^. The identification of pigments was conducted based on their distinctive retention times and absorption spectra (Zapata et al. 2000). Chl *a* and *β*-carotene were quantified according to Nagao et al. (2013), while zeaxanthin and *β*-cryptoxanthin were also quantified according to Nagao et al. (2023).

### Spectroscopies

Absorption spectra were measured at room temperature by a spectrophotometer (UV-2450; Shimadzu). Fluorescence spectra were measured at 77 K by a spectrofluorometer (RF-5300PC; Shimadzu).

## Results and discussion

### Protein compositions of PSI-LHCI, PSI, and LHCI

Three distinct preparations of PSI-LHCI, PSI, and LHCI, were purified from *G. sulphuraria* NIES-3638, and their trehalose density gradient centrifugation profiles (Figure 2) were similar to those of *C. caldarium* NIES-2137 (Nagao et al. 2023). SDS-PAGE analysis showed several bands, labeled a–o, in the three preparations (Figure 3). Mass spectrometry identified the major components of these bands as PSI and LHC proteins (Table 1). The bands b and m appeared slightly above the molecular weight marker of 29.0 kDa (Figure 3) and contained one LHC protein (Table 1). In contrast, the bands c and n appeared between the molecular weight markers of 20.1 and 14.3 kDa (Figure 3) and contained three LHC proteins (Table 1). The amino acid sequences of the four LHC proteins were similar to Lhcr1, Lhcr2, Lhcr4, and Lhcr5, as previously reported by Marquardt et al. (2000) and Marquardt et al. (2001). The accession IDs of the LHC proteins detected here were linked to the corresponding gene names, and these observations are summarized in Table 1.

**Figure 3.**
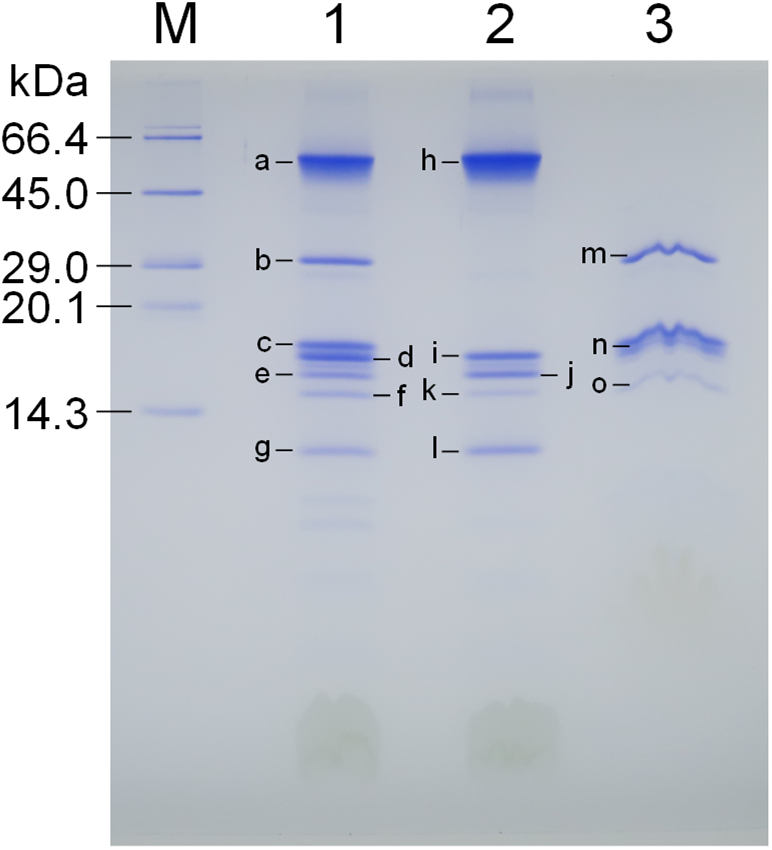
SDS-PAGE analysis of PSI-LHCI, PSI, and LHCI. Bands a–o were analyzed by mass spectrometry (see Table 1). Lane M, molecular marker; lane 1, PSI-LHCI; lane 2, PSI; lane 3, LHCI.

**Table 1.**
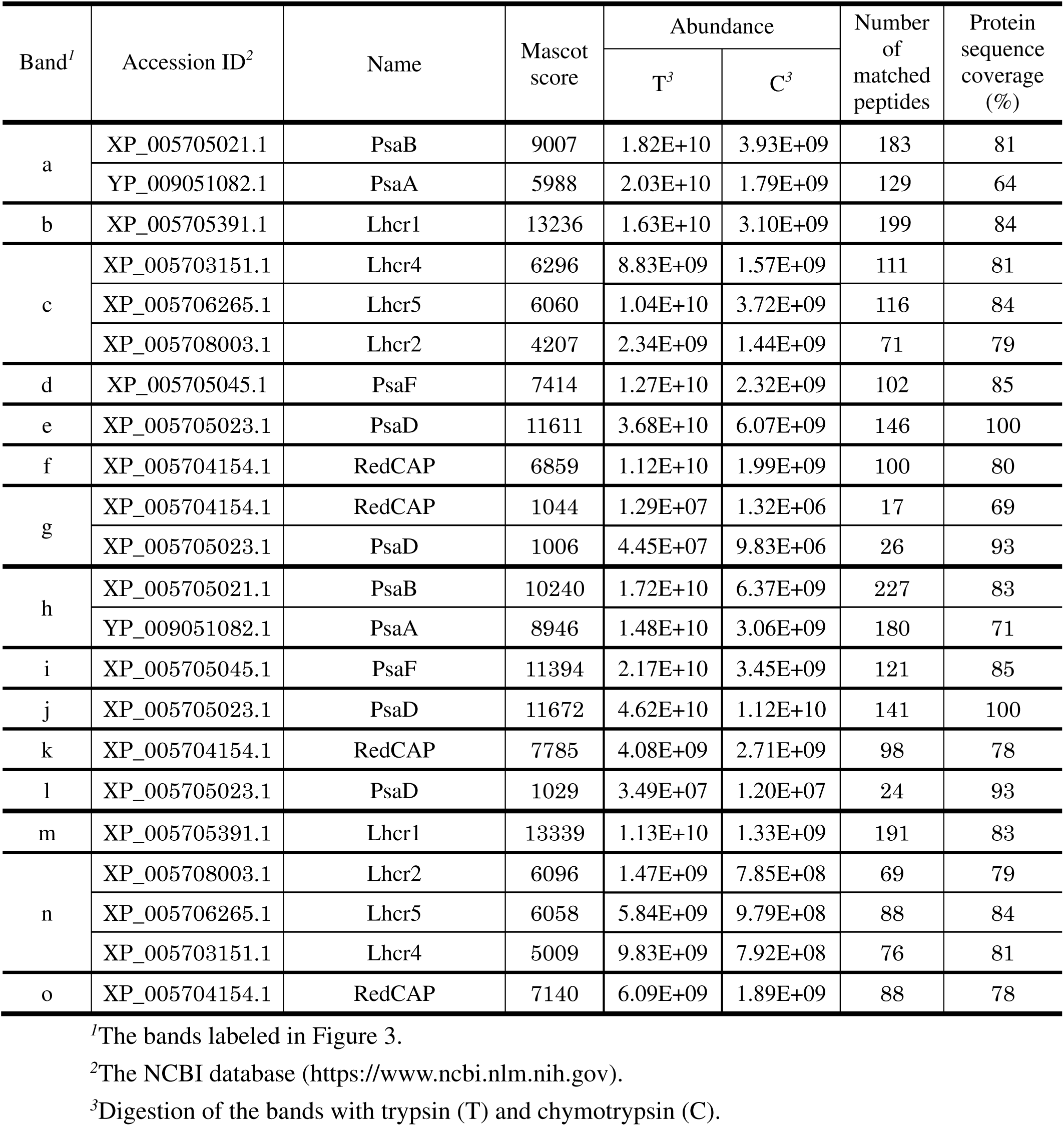
Major components in the SDS-PAGE bands by mass spectrometry.

| Band<sup>1</sup> | Accession ID<sup>2</sup> | Name | Mascot score | Abundance |  | Number of matched peptides | Protein sequence coverage (%) |
| --- | --- | --- | --- | --- | --- | --- | --- |
|  |  |  |  | T<sup>3</sup> | C<sup>3</sup> |  |  |
| a | XP_005705021.1 | PsaB | 9007 | 1.82E+10 | 3.93E+09 | 183 | 81 |
|  | YP_009051082.1 | PsaA | 5988 | 2.03E+10 | 1.79E+09 | 129 | 64 |
| b | XP_005705391.1 | Lhcr1 | 13236 | 1.63E+10 | 3.10E+09 | 199 | 84 |
| c | XP_005703151.1 | Lhcr4 | 6296 | 8.83E+09 | 1.57E+09 | 111 | 81 |
|  | XP_005706265.1 | Lhcr5 | 6060 | 1.04E+10 | 3.72E+09 | 116 | 84 |
|  | XP_005708003.1 | Lhcr2 | 4207 | 2.34E+09 | 1.44E+09 | 71 | 79 |
| d | XP_005705045.1 | PsaF | 7414 | 1.27E+10 | 2.32E+09 | 102 | 85 |
| e | XP_005705023.1 | PsaD | 11611 | 3.68E+10 | 6.07E+09 | 146 | 100 |
| f | XP_005704154.1 | RedCAP | 6859 | 1.12E+10 | 1.99E+09 | 100 | 80 |
| g | XP_005704154.1 | RedCAP | 1044 | 1.29E+07 | 1.32E+06 | 17 | 69 |
|  | XP_005705023.1 | PsaD | 1006 | 4.45E+07 | 9.83E+06 | 26 | 93 |
| h | XP_005705021.1 | PsaB | 10240 | 1.72E+10 | 6.37E+09 | 227 | 83 |
|  | YP_009051082.1 | PsaA | 8946 | 1.48E+10 | 3.06E+09 | 180 | 71 |
| i | XP_005705045.1 | PsaF | 11394 | 2.17E+10 | 3.45E+09 | 121 | 85 |
| j | XP_005705023.1 | PsaD | 11672 | 4.62E+10 | 1.12E+10 | 141 | 100 |
| k | XP_005704154.1 | RedCAP | 7785 | 4.08E+09 | 2.71E+09 | 98 | 78 |
| l | XP_005705023.1 | PsaD | 1029 | 3.49E+07 | 1.20E+07 | 24 | 93 |
| m | XP_005705391.1 | Lhcr1 | 13339 | 1.13E+10 | 1.33E+09 | 191 | 83 |
| n | XP_005708003.1 | Lhcr2 | 6096 | 1.47E+09 | 7.85E+08 | 69 | 79 |
|  | XP_005706265.1 | Lhcr5 | 6058 | 5.84E+09 | 9.79E+08 | 88 | 84 |
|  | XP_005703151.1 | Lhcr4 | 5009 | 9.83E+09 | 7.92E+08 | 76 | 81 |
| o | XP_005704154.1 | RedCAP | 7140 | 6.09E+09 | 1.89E+09 | 88 | 78 |

<sup>1</sup>The bands labeled in Figure 3.<sup>2</sup>The NCBI database (https://www.ncbi.nlm.nih.gov).<sup>3</sup>Digestion of the bands with trypsin (T) and chymotrypsin (C).

**Table 2.** Pigment compositions of PSI-LHCI, PSI, and LHCI.

| | zeaxanthin<sup>a</sup> | β-cryptoxanthin<sup>a</sup> | β-carotene<sup>a</sup> |
| --- | --- | --- | --- |
| PSI-LHCI | 134 ± 0 | 20 ± 2 | 145 ± 3 |
| PSI | 30 ± 0 | n.d.<sup>b</sup> | 165 ± 4 |
| LHCI | 322 ± 4 | 38 ± 0 | 118 ± 8 |

<sup>a</sup>Values represent millimole of pigment per mole of Chl *a* and are shown as means ± S.D. of three independent measurements.
<sup>b</sup>n.d., not detected.

The bands f, k, and o were observed between the molecular weight marker of 14.3 kDa and the bands c and n (Figure 3), corresponding to a hypothetical protein (Accession ID: XP_005704154.1) (Table 1). The amino acid sequence of XP_005704154.1 shares 61% similarity and 42% identity with that of g6493.t1 from the diatom *Chaetoceros gracilis* (Figure 4). The gene product of g6493.t1 has been identified as RedCAP, as reported in our recent study (Kato et al. 2024). RedCAPs are distinct from LHC proteins but are classified within the LHC protein superfamily (Engelken et al. 2010; Sturm et al. 2013). Therefore, the hypothetical protein (XP_005704154.1) from *G. sulphuraria* NIES-3638 is likely a RedCAP, as summarized in Table 1. Notably, RedCAP was also detected in band g, potentially due to partial proteolysis of this protein.

**Figure 4.**
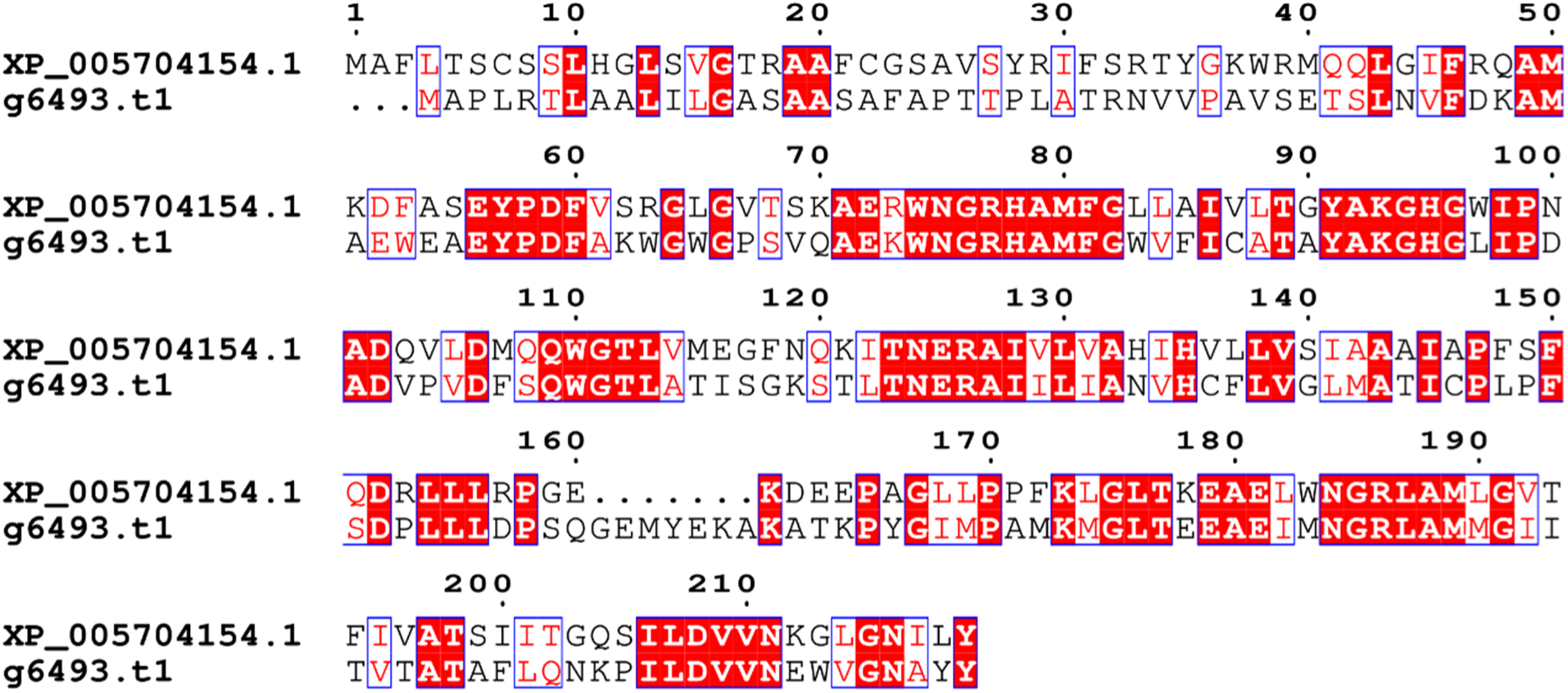
Comparison of the amino acid sequences of RedCAPs. Sequence alignment was performed using ClustalW (https://www.genome.jp/tools-bin/clustalw) and ESPript (https://espript.ibcp.fr/ESPript/cgi-bin/ESPript.cgi). XP_005704154.1, RedCAP of *G. sulphuraria*; g6493.t1, RedCAP of *C. gracilis*.

### Pigment compositions of PSI-LHCI, PSI, and LHCI

Figure 5 shows HPLC profiles of pigments extracted from the three preparations. PSI-LHCI and LHCI predominantly contained four types of pigments: zeaxanthin, *β*-cryptoxanthin, Chl *a*, and *β*-carotene (red and green lines, respectively), while PSI primarily contained three types of pigments: zeaxanthin, Chl *a*, and *β*-carotene (black line). The pigment stoichiometry was calculated as 134 for zeaxanthin, 20 for *β*-cryptoxanthin, and 145 for *β*-carotene in PSI-LHCI, and as 322, 38, and 118, respectively, in LHCI (Table 1). This indicates zeaxanthin enrichment in the *G. sulphuraria* LHCI. The Car composition of the *G. sulphuraria* LHCI resembled those of the *C. caldarium* LHCI (Nagao et al. 2023) and *C. merolae* LHCI (Tian et al. 2017).

**Figure 5.**
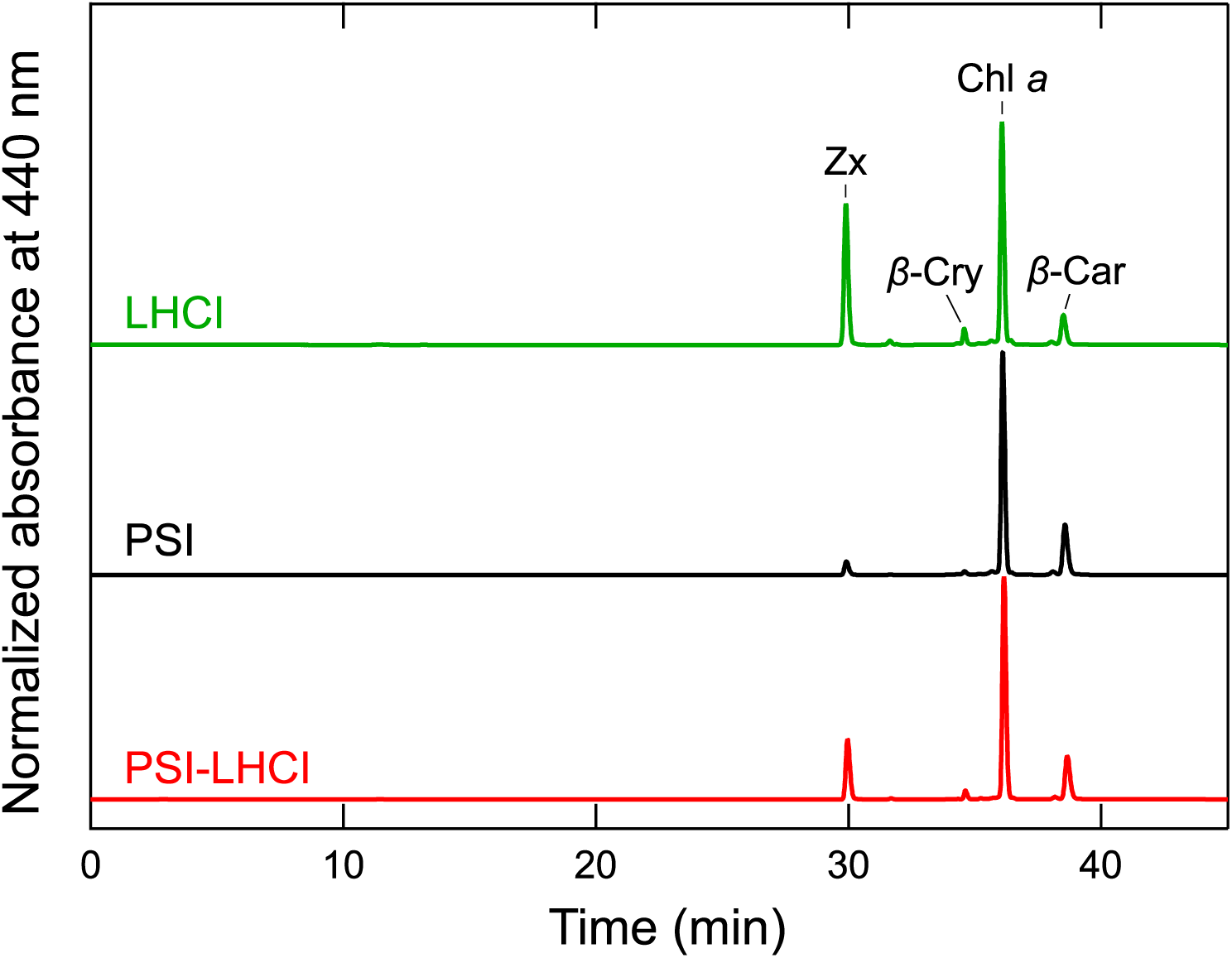
HPLC profiles of pigments extracts from PSI-LHCI, PSI, and LHCI. The elution profiles of the three preparations are indicated by red (PSI-LHCI), black (PSI), and green (LHCI) lines. The elution profiles were monitored at 440 nm and normalized by the peak intensity of Chl *a*. Zx, zeaxanthin; *β*-Cry, *β*-cryptoxanthin; Chl *a*, chlorophyll *a*; *β*-Car, *β*-carotene.

### Absorption and fluorescence-emission spectra of PSI-LHCI, PSI, and LHCI

Figure 6 shows absorption spectra of PSI-LHCI (red), PSI (black), and LHCI (green) measured at room temperature. The PSI-LHCI spectrum exhibited a Qy peak of Chl *a* at 679 nm, along with broad bands attributed to Chls *a* and Cars in the 400–500 nm range. The PSI spectrum showed a similar Qy peak of Chl *a* at approximately 679 nm, with corresponding bands in the 400–500 nm range. The LHCI spectrum displayed a Qy peak of Chl *a* at 670 nm, with the strongest intensities in the 400–500-nm bands. The Qy band of PSI-LHCI was broader than that of PSI, implying alterations in the energy levels of Chls by Chl-Chl interactions in interfaces between PSI and LHCIs and/or within LHCIs in PSI-LHCI.

**Figure 6.**
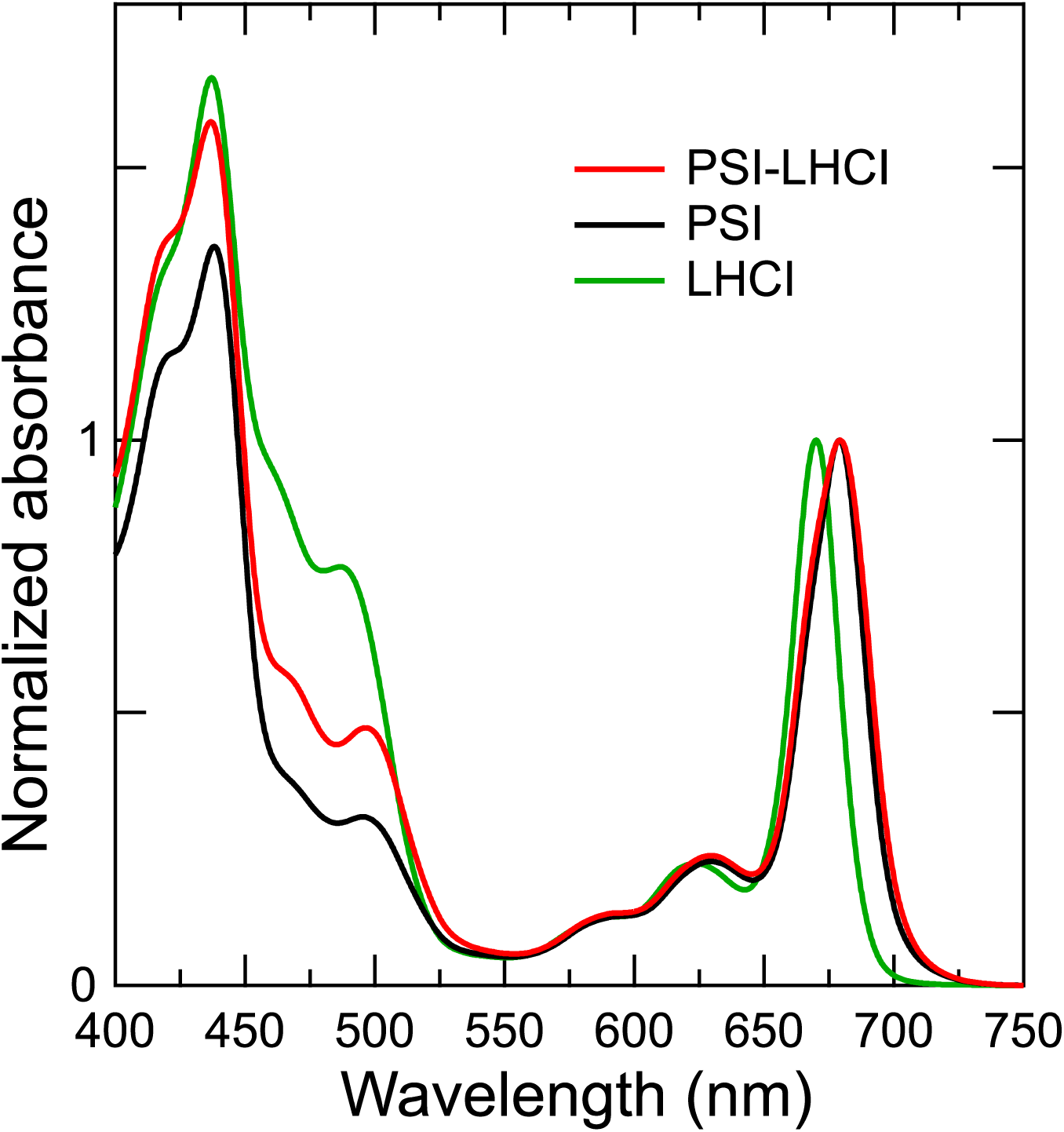
Absorption spectra of PSI-LHCI, PSI, and LHCI. The absorption spectra of the three preparations are indicated by red (PSI-LHCI), black (PSI), and green (LHCI) lines. The spectra were measured at room temperature and normalized by the intensity of the Chl-*a* Qy peak in each spectrum.

Figure 7 shows fluorescence spectra of PSI-LHCI (red), PSI (black), and LHCI (green) measured at 77 K. The PSI-LHCI spectrum depicted a fluorescence peak at 722 nm (red line), while the PSI spectrum displayed a peak at 721 nm and a small shoulder around 687 nm (black line). The LHCI spectrum exhibited a peak at 676 nm and a broad band around 733 nm (green line). The 722-nm peak in the PSI-LHCI spectrum was slightly red-shifted compared to the 721-nm peak in the PSI spectrum, with the 687-nm shoulder observed only in the PSI spectrum. The 721-nm peak likely originates from low-energy Chls within PSI, which have been well characterized in cyanobacterial PSIs (Kato et al. 2022; Schlodder et al. 2007; Toporik et al. 2020). In contrast, the LHCI spectrum showed little to no peaks at 721 and 722 nm, although it displayed a distinct vibronic band of Chl *a* at 733 nm (Niedzwiedzki and Blankenship 2010). The 722-nm peak in the PSI-LHCI spectrum may be attributed to structural changes in Chl environments by the binding LHCIs to PSI.

**Figure 7.**
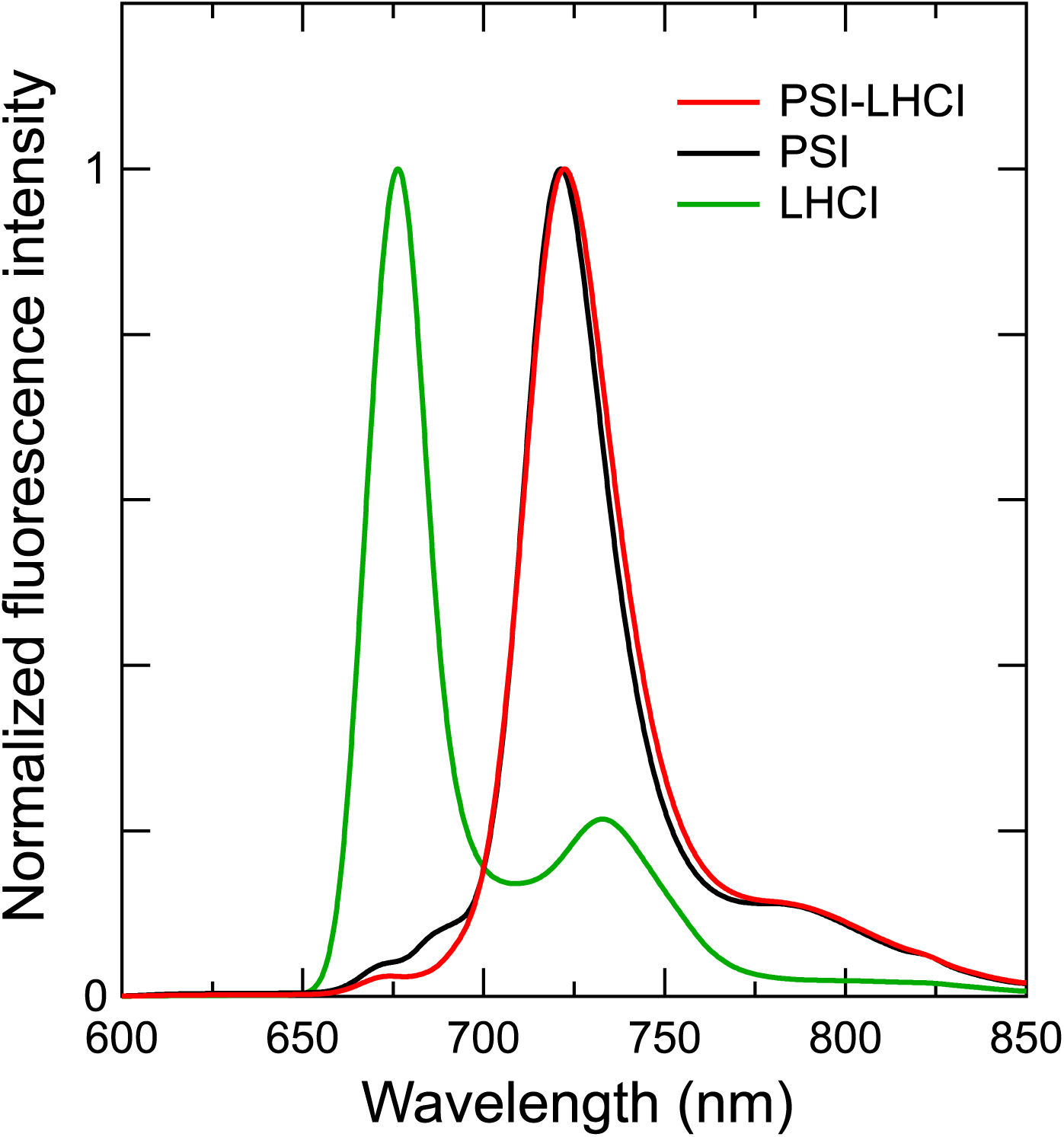
Fluorescence-emission spectra of PSI-LHCI, PSI, and LHCI. The fluorescence-emission spectra of the three preparations are indicated by red (PSI-LHCI), black (PSI), and green (LHCI) lines. The spectra were measured at 77 K upon excitation at 430 nm and normalized by the maximum-peak intensity in each spectrum.

### Purity of the *G. sulphuraria* PSI-LHCI, PSI, and LHCI

The polypeptide composition of PSI-LHCI comprised exclusively PSI and LHC proteins, with no detectable contamination from other proteins (Figure 3 and Table 1). The purity of PSI-LHCI was further corroborated by its fluorescence-emission spectrum, which exhibited little to no peak at 676 nm, attributed to LHCI (Figure 7). Similarly, LHCI was highly purified, as its polypeptide composition consisted solely of LHC proteins (Figure 3 and Table 1), and its fluorescence spectrum showed no emission at 722 nm, corresponding to PSI (Figure 7). Additionally, the Qy peak in the absorption spectrum of LHCI was notably blue-shifted relative to that of PSI-LHCI (Figure 6).

Unlike the two protein complexes, PSI exhibited low purity due to the presence of the polypeptide band of RedCAP (Figure 3 and Table 1). Although Anzergent 3-16 appears to harshly dissociate LHCI from PSI-LHCI, RedCAP persists in the PSI fraction (Figure 3 and Table 1), suggesting a strong interaction between RedCAP and PSI. Furthermore, as discussed in our recent study (Kato et al. 2024), the RedCAP subunit has been identified in nearly identical positions in the PSI-LHCI structures of *P. cruentum*, *Chaetoceros gracilis*, and *Chroomonas placoidea*, though it is absent in the PSI-LHCI structures of *C. caldarium* and *C. merolae*. Determining whether the binding properties of RedCAP in the PSI-LHCI of *G. sulphuraria* are conserved across red-lineage algae remains a key question.

### LHC protein superfamily in *G. sulphuraria*

SDS-PAGE analysis and mass spectrometry clearly identified the bands b and m as the unique LHC protein, Lhcr1, whose apparent molecular weight was significantly higher than those of other Lhcr proteins detected in the bands c and n (Figure 3 and Table 1). It is known that at least five *Lhcr* genes have been identified in *G. sulphuraria* (Marquardt et al. 2001; Marquardt et al. 2000), and LHC-related proteins of *G. sulphuraria* were immuno-detected in the molecular-weight range of 16–20 kDa (Marquardt et al. 2001). As Lhcr1 was observed near a molecular-weight marker of 29.0 kDa (Figure 3), the polypeptide bands of LHCs in *G. sulphuraria* appear to differ between the present study and the previous studies (Marquardt et al. 2001; Marquardt and Rhiel 1997). These differences may be attributable to variations in the preparation conditions of PSI-LHCI, as well as differences in the strains used, namely NIES-3638 in the present study versus SAG 107.79 in the previous studies (Marquardt et al. 2001; Marquardt and Rhiel 1997).

LHC proteins associated with PSI and PSII share a similar structure comprising three membrane-spanning helices, regardless of the species of the photosynthetic organisms (Green and Durnford 1996; Green and Pichersky 1994; Hippler and Nelson 2021; Shen 2022). Marquardt et al. (2001) showed that in *G. sulphuraria*, Lhcr1 has an N-terminal extension compared with Lhcr2–5. This N-terminal extension was distinctly observed in the present study, as the apparent molecular weight of Lhcr1 on SDS-PAGE was approximately 29.0 kDa (Figure 3). Given that the three helices in Lhcr1–5 of *G. sulphuraria* are thought to be conserved (Marquardt et al. 2001), the structure of the N-terminal region of Lhcr1 might play a role in its association with PSI and in the assembly of PSI-LHCI supercomplexes.

The *G. sulphuraria* PSI-LHCI supercomplex in the present study contained Lhcr1, Lhcr2, Lhcr4, Lhcr5, and RedCAP within the LHC protein superfamily (Figure 3 and Table 1). However, Lhcr3, identified by Marquardt et al. (2001), was not observed as a major component. This absence may be because of (i) no expression of Lhcr3 in the NIES-3638 strain or (ii) removal of Lhcr3 during preparation of PSI-LHCI. Marquardt et al. (2001) also reported that the *G. sulphuraria* PSI-LHCI likely comprises 7–9 LHCI subunits and a PSI-monomer core, based on negative-stain electron microscopy. Among red algae, structural studies of PSI-LHCI revealed five LHCI subunits in *C. caldarium* (Kato et al. 2024), three to five in *C. merolae* (Antoshvili et al. 2019; Pi et al. 2018), and eight in *P. cruentum* (You et al. 2023). RedCAP was one of the eight LHCI subunits found in the *P. cruentum* PSI-LHCI structure (You et al. 2023), whereas it was absent in the PSI-LHCI structures of *C. caldarium* (Kato et al. 2024) and *C. merolae* (Antoshvili et al. 2019; Pi et al. 2018). Because the *G. sulphuraria* PSI-LHCI purified here contains a RedCAP in addition to LHC proteins (Figure 3 and Table 1), its overall structure and light-harvesting strategy may be similar to those of the *P. cruentum* PSI-LHCI, rather than the PSI-LHCI of *C. caldarium* or *C. merolae*. To address these differences, further structural and functional studies of the *G. sulphuraria* PSI-LHCI are needed to better understand the diversity of LHC proteins and RedCAP in red-lineage algal PSI-LHCI supercomplexes.

## Acknowledgements

We thank Kumiyo Kato and Satoko Kakiuchi for their assistance in this study. This work was supported by JSPS KAKENHI grant Nos. JP23K14211 (Y.N.), JP22H04916 (J.-R.S.), and JP23H02423 (R.N.), and Takeda Science Foundation (K.K.).

## Author Contributions

R.N. conceived the project; R.N. and H.O. performed isolation and characterization of PSI-LHCI, PSI, and LHCI; T.S. and N.D. performed mass spectrometry; K.K and Y.N. provided laboratory resources and supervised experimental work; and R.N. drafted the original manuscript; J.-R.S. modified the manuscript; and R.N. wrote the final manuscript, and all authors joined the discussion of the results.

